# A significant enrichment that is not: spatial nulls, co-expression, and the imaging transcriptomics of EEG alpha-power genetics

**DOI:** 10.64898/2026.07.23.740331

**Authors:** Jacopo Schenetti

## Abstract

**Background:** Imaging transcriptomics routinely asks whether a trait-associated gene set is over-expressed in a region of interest, and the field standard is to guard that inference with a spatial-autocorrelation-preserving spin test. Electroencephalographic (EEG) oscillatory power is among the most heritable human neurophysiological traits, and the cortical generators of the alpha rhythm have been characterised independently from resting-state magnetoencephalography. That makes it a natural test bed for two questions at once: whether trait genetics is regionally organised, and what such a test actually establishes.

**Objective:** To test whether alpha-associated genetic signal is spatially enriched in the cortical generators of the alpha rhythm, and to evaluate that inference against complementary null models.

**Methods:** MAGMA gene-based analysis of ENIGMA-EEG summary statistics for six phenotypes: central and occipital alpha power, occipital alpha peak frequency, and theta, beta and delta power. Regional transcription was obtained from the Allen Human Brain Atlas (AHBA) with *abagen* in the Glasser HCP-MMP1.0 atlas, with Schaefer-100 and Yan-600 as sensitivity analyses. Enrichment in the 41 cortical alpha-source regions was quantified with a threshold-free continuous score and a top-100 gene-set composite, and assessed against three complementary nulls: a spin test (10,000 rotations, cross-checked against brainsmash surrogates), a co-expression-aware gene-set null (10,000 matched random gene sets), and a positive control on a known expression gradient.

**Results:** Judged by the field-standard spin test alone, this study would have reported a positive, biologically coherent finding: alpha-power genes are enriched in the cortical alpha generators (continuous *p*_spin = 0.022; top-100 *p*_spin = 0.030), with the spin result corroborated by an independent surrogate model (*p* = 0.018) and the pipeline validated by a positive control (*p*_spin = 2 × 10^−4^). Three further tests undo that conclusion. First, it is not band-specific: theta, beta and delta enrich comparably or more strongly (top-100 beta *p*_spin = 0.011; delta 0.042; continuous theta 0.042). Second, it does not replicate across alpha phenotypes (occipital alpha continuous *p*_spin = 0.20; alpha peak frequency non-significant, *p*_spin ≥ 0.066). Third, against random gene sets of matched size the alpha set is unremarkable (*p*_geneset = 0.33): the apparent enrichment is a generic property of arbitrary gene sets in this cortical territory, and a spatial-only null cannot see it. No test survived false-discovery-rate correction across the 24-cell phenotype × score × region-set grid (minimum *q* = 0.127), and nominal significance did not survive a change of parcellation.

**Conclusion:** EEG alpha-power genetics shows no regionally specific transcriptomic signature in the cortical generators of the rhythm; the weak tendency that is present is shared across frequency bands, consistent with their known genetic correlation. Methodologically, it shows that correcting for spatial autocorrelation is necessary but not, on its own, enough: an enrichment that passes the spin test, is backed by a surrogate model, and reads as mechanistically plausible can still be explained entirely by gene-set co-expression. Enrichment claims in imaging transcriptomics should report a gene-set null alongside the spatial null.

## 1. Introduction

The occipito-parietal alpha rhythm (∼8–13 Hz) is the dominant oscillation of the awake human EEG at rest. Twin studies place the heritability of alpha power and peak frequency among the highest of any neurophysiological trait (h^2^ ≈ 0.7–0.8; Smit et al., 2005), and a meta-analytic GWAS of EEG oscillatory power by the ENIGMA-EEG working group (Smit et al., 2018) identified common variants associated with band-limited power across delta, theta, alpha and beta, linking several psychiatric-liability genes to oscillatory brain activity.

Where in the cortex that heritable variance acts is a separate question. The alpha rhythm is not generated uniformly: Tabarelli et al. (2022) characterised its cortical generators directly, using resting-state magnetoencephalography (MEG) from the Cam-CAN cohort and minimum-norm source reconstruction in the Glasser HCP-MMP1.0 multimodal parcellation to identify 41 cortical alpha-rhythm sources spanning occipital, parietal and frontal sectors. If the genetic architecture of alpha power were regionally organised, so that the genes shaping how much alpha a person produces acted preferentially where that alpha is generated, those 41 regions should carry a distinguishable molecular signature. The question has practical as well as basic interest, since brain-state-dependent EEG–TMS protocols read out and target the same generators (Zrenner et al., 2018), yet it has never been tested.

Imaging transcriptomics provides the natural test. By correlating macroscale brain maps with regional gene-expression profiles from the Allen Human Brain Atlas (AHBA; Hawrylycz et al., 2012) via toolboxes such as *abagen* (Markello et al., 2021), it has become a mature approach for grounding neuroimaging phenotypes in molecular architecture (Arnatkeviciute et al., 2019; Fornito et al., 2019). It has not, however, been applied to an EEG-trait GWAS.

That gap motivates a second, methodological question. Enrichment tests of this kind are conventionally guarded with a spin test, which permutes brain maps while preserving their spatial autocorrelation (Alexander-Bloch et al., 2018). But spatial autocorrelation is not the only structure that can manufacture an apparent enrichment: genes within a set are themselves spatially co-expressed, which inflates significance in a way a spatial-only null cannot see (Fulcher et al., 2021). How much difference this makes in practice, on a real trait, with the outcome not known in advance, is rarely demonstrated.

We therefore address two questions together:

**H1 (substantive)**. Genes associated with EEG alpha power are expressed more highly in the cortical alpha generators (Tabarelli et al., 2022) than in the rest of cortex, and this enrichment is *specific* to the alpha band.

**H2 (methodological)**. Whatever H1’s outcome under a spin test, does that verdict survive complementary null models — a co-expression-aware gene-set null, a positive control, and an independent surrogate model?

The design is built to yield an informative answer even from a modestly powered GWAS. *Specificity* is assessed with an **internally power-matched** comparison: three non-alpha bands and two further alpha phenotypes pass through the identical pipeline, so the relative contrast between them is meaningful regardless of the absolute power of any one GWAS. *Sensitivity* is established with a positive control the test must recover. *Robustness* is examined across three parcellations of different granularity. As we report below, the spin test alone supports H1; the remaining tests then take it apart, with implications that reach beyond this particular trait.

## 2. Methods

All analyses used publicly available secondary data. No primary data were collected. Code and derived results are available (see *Data and code availability*); raw GWAS, AHBA and reference-panel data are redistributed under their original licences only as download instructions.

### 2.1 GWAS summary statistics

We obtained ENIGMA-EEG oscillatory-power summary statistics (Smit et al., 2018; 2018-11-01 release, select_N6000_Hom100) for six scalp-derived phenotypes: alpha power at the central electrode Cz (**alphaCz**, the pre-declared primary phenotype), alpha power at an occipital site (**alphaOcc**), the occipital alpha peak frequency (**peakOcc**, an index of individual alpha frequency), and, as specificity controls, theta, beta and delta power at Cz (**thetaCz, betaCz, deltaCz**). Each file contained ≈4.96 million autosomal SNPs with per-SNP sample size (N ≈ 6,000–7,700). The genome build was verified as GRCh37/hg19 (e.g., rs3094315 at chr1:752,566), matching the gene-location and reference-panel builds, so no liftOver was required.

### 2.2 Gene-based analysis (MAGMA)

Gene-based association was computed with MAGMA v1.10 (de Leeuw et al., 2015). SNPs were annotated to genes (build-37 NCBI gene locations; 19,427 genes) with a 35 kb upstream / 10 kb downstream window; the 18,053 genes with ≥1 mapped SNP were carried into the gene-based test. The SNP-wise mean model used the 1000 Genomes European reference panel (1000 Genomes Project Consortium, 2015; 503 individuals) to model linkage disequilibrium. Entrez gene identifiers were mapped to HGNC symbols using the gene-location file. Gene *p*-values were corrected by Bonferroni and Benjamini–Hochberg FDR (Benjamini & Hochberg, 1995). For the enrichment analysis we retained (i) the full genome-wide vector of gene-level *z*-statistics and (ii) a top-100 gene set ranked by gene *p*-value; threshold-based sets (Bonferroni, FDR, *p*<10^−3^, *p*<10^−2^) were also generated for reference.

### 2.3 Regional gene expression (AHBA / abagen)

Regional transcription was estimated with *abagen* (development build post-0.1.3, GitHub `netneurolab/abagen`, required for pandas ≥2 compatibility; Markello et al., 2021) from AHBA microarray data (Hawrylycz et al., 2012). Default processing was used (intensity-based filtering threshold 0.5; probe selection by differential stability (Hawrylycz et al., 2015); within-donor scaled robust sigmoid normalisation; `missing=“interpolate”;` matched-sample normalisation). Of the six AHBA donors, five were used; donor 15496 (H0351.1015, a left-hemisphere-only donor) was excluded because its normalized-microarray archive was unavailable at source at the time of analysis (persistent HTTP 404). The primary parcellation was Glasser HCP-MMP1.0 (Glasser et al., 2016; 360 regions), matching the space in which the alpha sources are defined. Because AHBA samples are mapped by *abagen* to fsaverage5, the Glasser atlas was resampled from fsLR32k to fsaverage5 using the Human Connectome Project `resample_fsaverage` registration spheres (Van Essen et al., 2012; Glasser et al., 2013; nearest-neighbour in the common fsaverage frame); resampling preserved all 180 parcels per hemisphere. The resulting expression matrix contained 360 regions × 15,563 genes, with complete coverage of all 82 target parcels. For robustness, expression was also computed in the Schaefer-100 (Schaefer et al., 2018) and Yan-600 homotopic (Yan et al., 2023) parcellations, both in fsaverage5.

### 2.4 Alpha-source region set

The 41 alpha-source regions of Tabarelli et al. (2022) were used as the target set (16 occipital, 7 parietal, 18 frontal HCP-MMP labels), applied bilaterally (82 parcels). Because the sources are natively defined in the Glasser atlas, the mapping is one-to-one by label, avoiding subjective region translation. A prefrontal-hub subset (areas 8C, 8Av, 9a; 6 parcels) was additionally tested, motivated by the paper’s identification of 8C as the principal connectivity hub. For the Schaefer-100 and Yan-600 sensitivity analyses, the alpha-source set was re-mapped by fsaverage5 surface overlap: a parcel was flagged as target if ≥50% of its cortical vertices fell within the Glasser alpha-source set (31/100 Schaefer parcels; 180/600 Yan parcels).

### 2.5 Alpha transcriptomic score and enrichment test

We defined two per-region scores. The **continuous** score is the Spearman correlation, across all genes shared between the expression matrix and the gene-based results, between a region’s expression profile and the genome-wide gene-level *z*-statistics; it uses the full association signal without any significance threshold. The **top-100** score is the mean *z*-scored expression, across regions, of the top-100 gene set. Enrichment was quantified as the difference between the mean score in the target regions and the rest of cortex.

Statistical significance was assessed against a spin-test null that preserves the spatial autocorrelation of cortical maps (Alexander-Bloch et al., 2018). Because the parcellation was surface-based, we generated 10,000 spatial rotations of the parcel centroids on the fsaverage5 sphere (nearest-neighbour reassignment; left- and right-hemisphere rotations mirror-matched; seed fixed and logged), and computed a one-sided *p*-value (H1: target > rest) as the fraction of rotated maps with a target-region mean at least as large as observed. This self-contained implementation was used because the spin generator had been removed from the version of *netneurotools* (Liu et al., 2025) available at analysis time. Specificity and replication were assessed by repeating the test for all six phenotypes and both scores, and correcting the resulting 24-cell grid by Benjamini–Hochberg FDR. Robustness was assessed by repeating the primary-phenotype test in the Schaefer-100 and Yan-600 parcellations. Because the analysis is internally power-matched (every phenotype shares the same GWAS design, the same expression matrix and the same null), the *relative* comparison between the alpha and control bands is informative about specificity independent of the absolute statistical power of any single GWAS.

### 2.6 Validation and complementary nulls

Three analyses established the validity of the enrichment test. **(i) Positive control**. To confirm that the pipeline can detect a genuine spatial effect, we replaced the gene statistic with the gene loadings of the first principal component (PC1) of the AHBA expression matrix, the dominant and well-replicated cortical expression gradient, and set the target to the top-tertile PC1 regions; a correct pipeline must recover a highly significant enrichment. **(ii) Co-expression-aware geneset null**. Because gene-set enrichment on AHBA is prone to false positives from within-set spatial co-expression (Fulcher et al., 2021), we complemented the spin test (which permutes space) with a gene-set null for the top-100 analysis: 10,000 random gene sets of matched size were drawn, and *p*_geneset was the fraction whose target-versus-rest difference met or exceeded the observed value. **(iii) Spin validation**. Because the spin generator was self-implemented, we benchmarked it against an independent spatial-autocorrelation-preserving surrogate model (brainsmash; Burt et al., 2020; Markello & Mišić, 2021), generating 1,000 surrogate maps of the primary alpha score over a Euclidean parcel-distance matrix and comparing the resulting *p*-value with the spin *p*-value.

### 2.7 Reproducibility

Every stage logged its parameters (random seed, parcellation version, *abagen* and package versions, MAGMA build) to a machine-readable run log. A global seed (20260721) governed all permutations. Analyses ran under Python 3.12.

## 3. Results

### 3.1 Gene-based analysis recovers credible signal

MAGMA gene-based analysis of central alpha power tested 18,053 genes. A single gene survived genome-wide Bonferroni and FDR correction, *PRKG2* (*p* = 4.0 × 10^−7^, *q* = 0.007), consistent with the modest sample size. The leading sub-threshold signal included the chr3p21 pleiotropic cluster (*GNL3, GLT8D1, NT5DC2, PBRM1*; *p* ≈ 4–5 × 10^−5^), one of the most robust loci in psychiatric genetics and consistent with the ENIGMA-EEG thesis that psychiatric-liability genes contribute to oscillatory brain activity (Smit et al., 2018). The presence of this expected locus provides a positive control that the gene-based pipeline behaves correctly. Because only one gene survived strict correction, we did not treat a threshold-based gene set as the primary readout; instead the continuous score (using the full gene-level signal) was pre-declared primary and the top-100 gene set served as a sensitivity analysis. (The analysis grid was fixed in advance but was not formally pre-registered.)

### 3.2 The enrichment test is valid and the custom spin is corroborated

The positive control confirmed that the pipeline detects genuine spatial structure: with the PC1 expression-gradient loadings as the gene statistic and the top-tertile PC1 regions as the target, the spin test recovered a large, highly significant enrichment (target-versus-rest difference = 0.74; *p*_spin = 2 × 10^−4^). The subsequent alpha nulls are therefore not a floor effect of an insensitive method. The self-implemented spin was corroborated by an independent surrogate model: for the primary alpha map, the spin *p*-value (0.022) and the brainsmash *p*-value (0.018) were closely concordant.

### 3.3 By the spin test alone, alpha genes are enriched in the alpha generators

In the primary Glasser parcellation, central alpha power showed enrichment of alpha transcriptomic signal in the 41 alpha-source regions for both scores: the continuous score gave a target-versus-rest difference corresponding to *z* = 1.98 relative to the spin null (*p*_spin = 0.022), and the top-100 score gave *z* = 1.77 (*p*_spin = 0.030). The prefrontal-hub subset (8C, 8Av, 9a) was not significant for either score (continuous *p*_spin = 0.11; top-100 *p*_spin = 0.73), indicating that any enrichment was distributed across the broad alpha-source network rather than concentrated at the connectivity hubs.

This is the point at which a conventional analysis would stop. Two independent scores agree, the spatial null is the field standard, an independent surrogate model reproduces the *p*-value, the pipeline has passed a positive control, and the result has a ready mechanistic reading: the genes that shape alpha power are over-expressed where alpha is generated. H1 appears supported. The three analyses that follow show that it is not.

### 3.4 First test: the enrichment is not band-specific and does not replicate

Repeating the test across all six phenotypes overturned the primary interpretation (Table 1). In the continuous score, the theta control band was also nominally enriched in the alpha-source regions (*p*_spin = 0.042), while the occipital alpha phenotype, the natural replication of the central alpha result, was not (*p*_spin = 0.20). In the top-100 score, the pattern was still less specific: the occipital alpha phenotype (*p*_spin = 0.008) and the beta control band (*p*_spin = 0.011) gave the strongest signals of all, both exceeding central alpha (*p*_spin = 0.030), with delta also nominally significant (*p*_spin = 0.042). The alpha peak-frequency (IAF) phenotype, arguably the most closed-loop-relevant since the controller locks to it, was not significant for either score (continuous *p*_spin = 0.13; top-100 *p*_spin = 0.066). After Benjamini–Hochberg correction across the 24-cell phenotype × score × region-set grid, **no test remained significant** (minimum *q* = 0.127). The prefrontal-hub subset was null or negative throughout.

**Table 1.** Spin-test enrichment of alpha transcriptomic signal in the 41 alpha-source regions (Glasser HCP-MMP1.0), by phenotype and score. *z* is relative to the 10,000-rotation spin null; *p*_spin is one-sided (target > rest); *q* is Benjamini–Hochberg FDR across the full 24-cell (6 phenotype × 2 score × 2 region-set) grid. Band labels distinguish alpha power, the alpha peak-frequency (IAF) phenotype, and the non-alpha control bands.

| Score | Phenotype | Band | $z$ vs null | $p_{\text{spin}}$ | $q$ (FDR) |
| --- | --- | --- | --- | --- | --- |
| Continuous | alphaCz | alpha-power | 1.98 | 0.022 | 0.169 |
| Continuous | alphaOcc | alpha-power | 0.87 | 0.203 | 0.325 |
| Continuous | peakOcc | alpha-frequency | 1.16 | 0.130 | 0.283 |
| Continuous | thetaCz | control | 1.68 | 0.042 | 0.169 |
| Continuous | betaCz | control | 1.22 | 0.123 | 0.283 |
| Continuous | deltaCz | control | 0.99 | 0.168 | 0.310 |
| Top-100 | alphaCz | alpha-power | 1.77 | 0.030 | 0.169 |
| Top-100 | alphaOcc | alpha-power | 2.28 | 0.008 | 0.127 |
| Top-100 | peakOcc | alpha-frequency | 1.72 | 0.066 | 0.227 |
| Top-100 | betaCz | control | 2.13 | 0.011 | 0.127 |
| Top-100 | deltaCz | control | 1.62 | 0.042 | 0.169 |
| Top-100 | thetaCz | control | 1.32 | 0.079 | 0.237 |

### 3.5 Second test: the alpha gene set is indistinguishable from random gene sets

This test replaces the spatial null with a gene-set null. Assessed against 10,000 random gene sets of the same size, rather than against rotated brain maps, the observed target-versus-rest difference (0.060) proved unremarkable: it fell near the centre of the random-gene-set distribution (mean 0.043; *p*_geneset = 0.33; Fig. 4B). Arbitrary gene sets are, on average, mildly over-expressed in this occipito-parieto-frontal territory, and the alpha set is not distinguishable from them.

**Figure 1:**
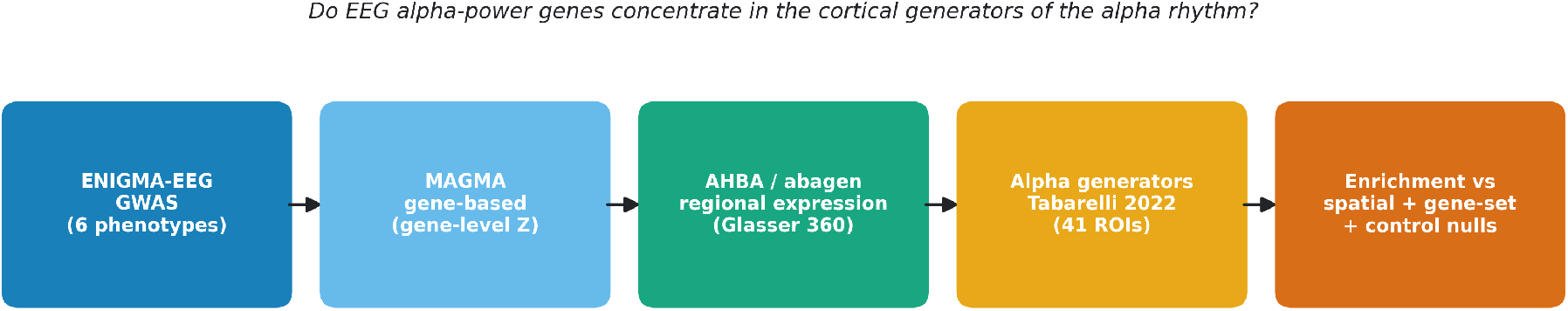
Study design. From the ENIGMA-EEG oscillatory-power GWAS through MAGMA gene-based analysis and AHBA regional expression to the cortical alpha-source ROIs, tested for spatial enrichment against complementary null models.

**Figure 2:**
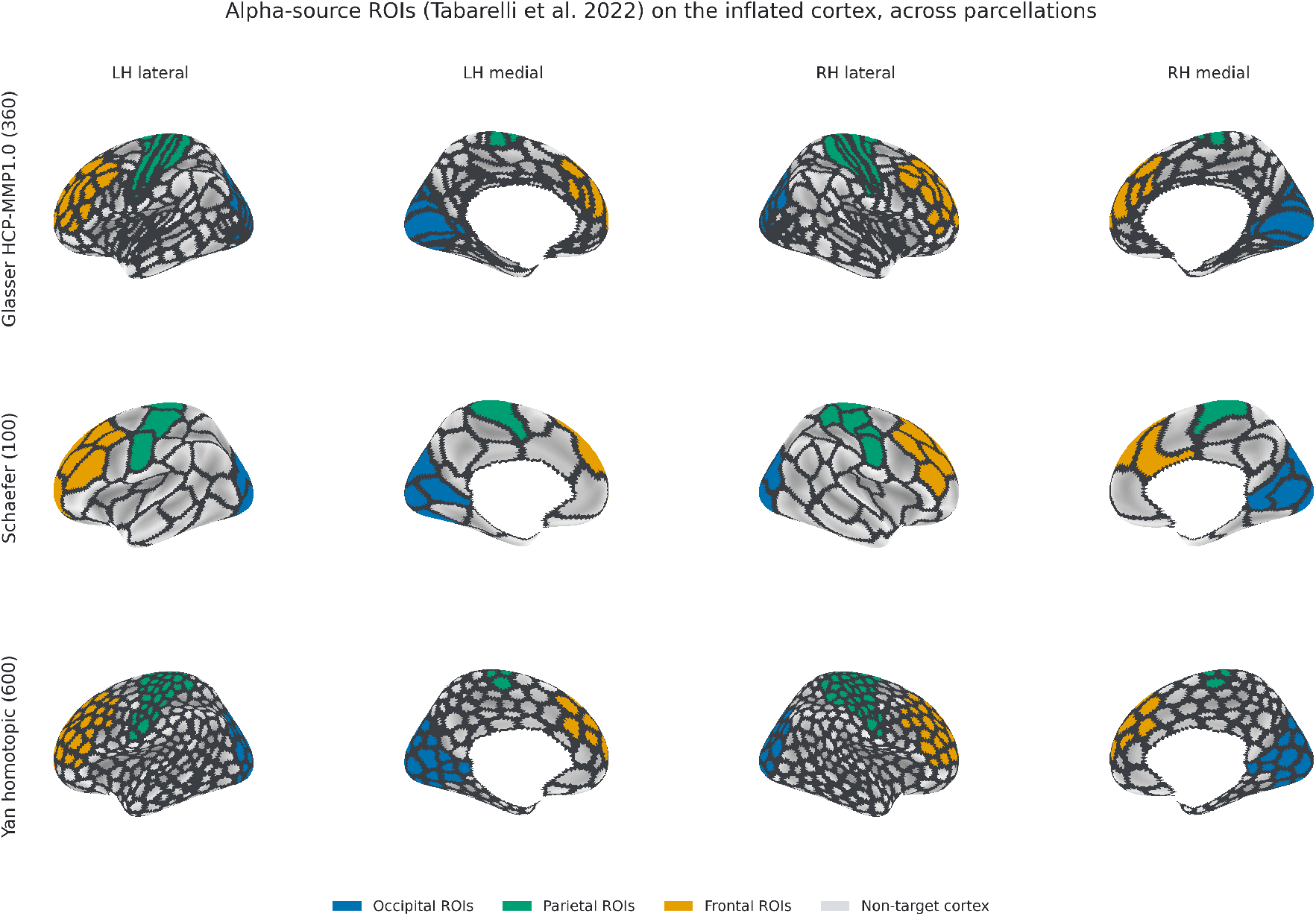
Alpha-source ROIs on the inflated cortex, across parcellations. The 41 Tabarelli alpha-source regions (bilateral) shown on the fsaverage5 inflated surface for Glasser HCP-MMP1.0, Schaefer-100 and Yan-600. In every parcellation the target parcels are coloured by sector (occipital, parietal, frontal); for Schaefer and Yan each target parcel takes the sector of the Glasser alpha-source it most overlaps. Thin dark lines are atlas parcel boundaries; grey is non-target cortex.

**Figure 3:**
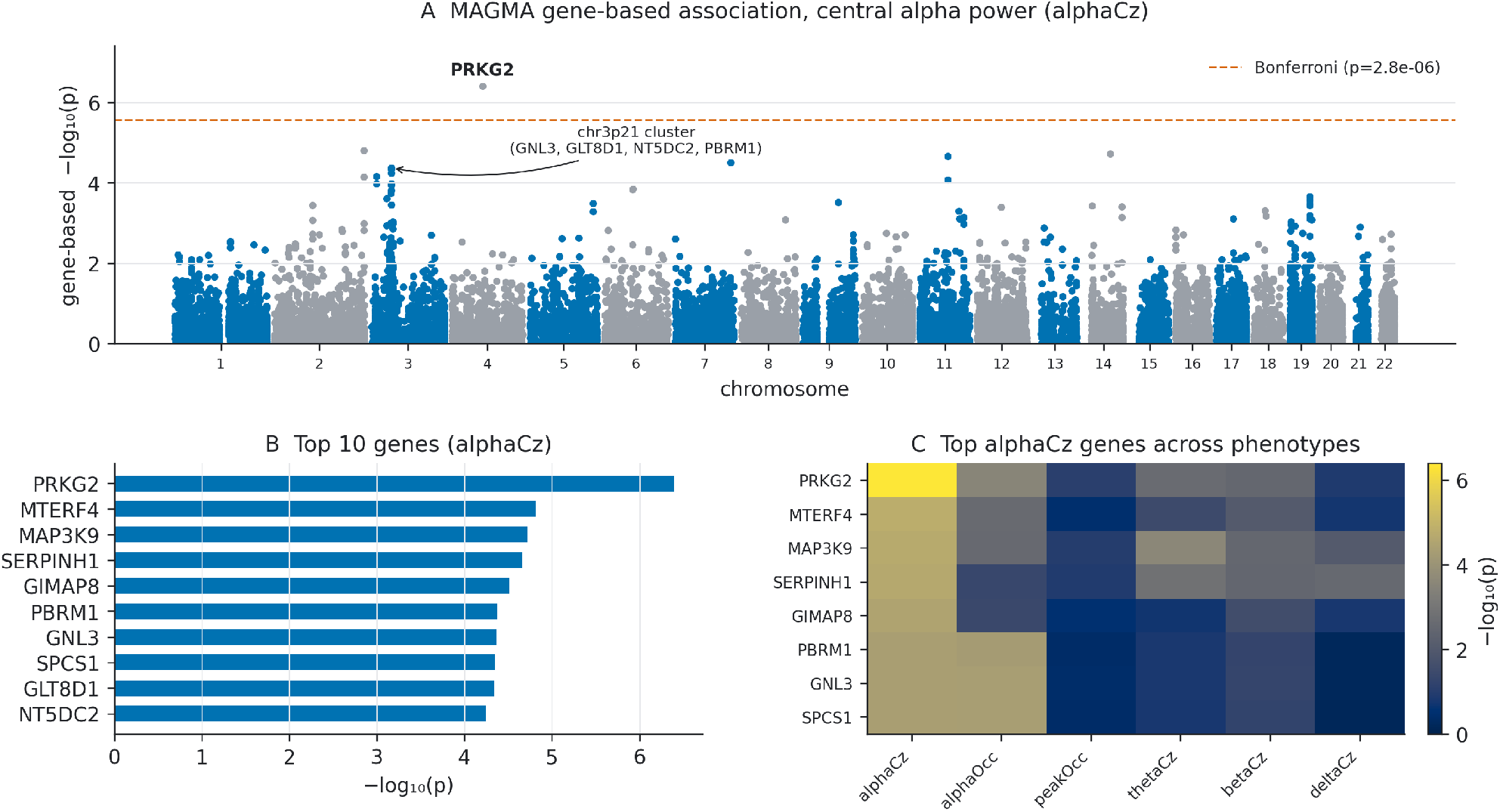
Gene-based genetics. (A) MAGMA gene-based association for central alpha power (alphaCz); *PRKG2* passes Bonferroni and the chr3p21 pleiotropic cluster is the leading sub-threshold signal. (B) Top-10 genes. (C) −log_10_ *p* of the top alphaCz genes across all six phenotypes — the gene-level signal is largely specific to alphaCz.

**Figure 4:**
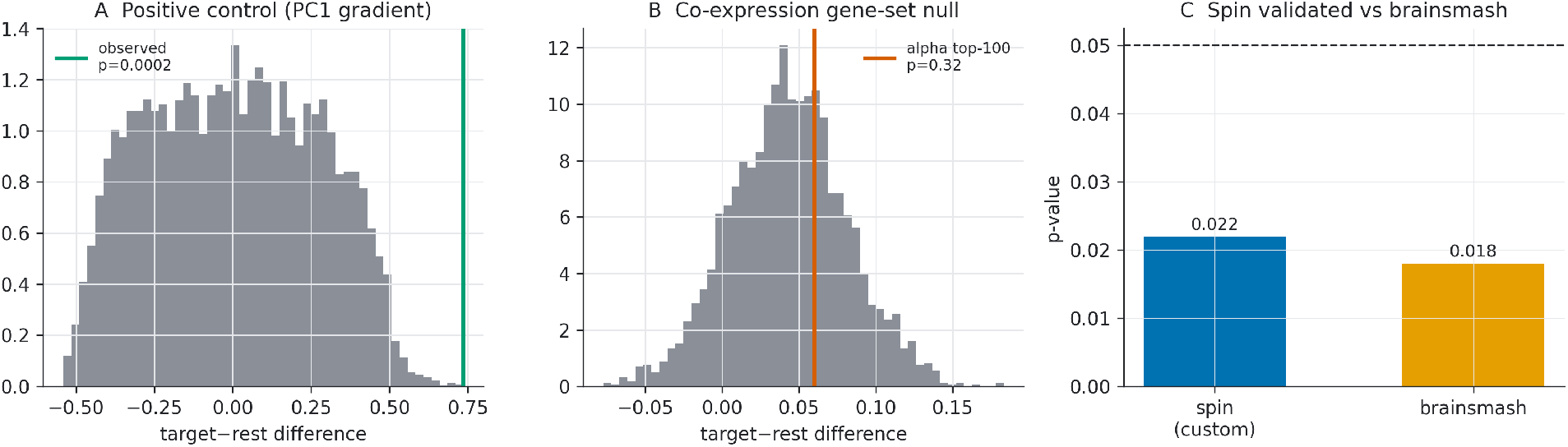
Validation of the enrichment test. (A) Positive control: with the dominant expression gradient (PC1) as the gene statistic, the spin test recovers a large enrichment (*p*_spin = 2 × 10^−4^). (B) Co-expression-aware gene-set null: the alpha top-100 target-versus-rest difference sits at the centre of 10,000 random gene sets (*p*_geneset = 0.33). (C) The custom spin agrees with brainsmash surrogates.

**Figure 5:**
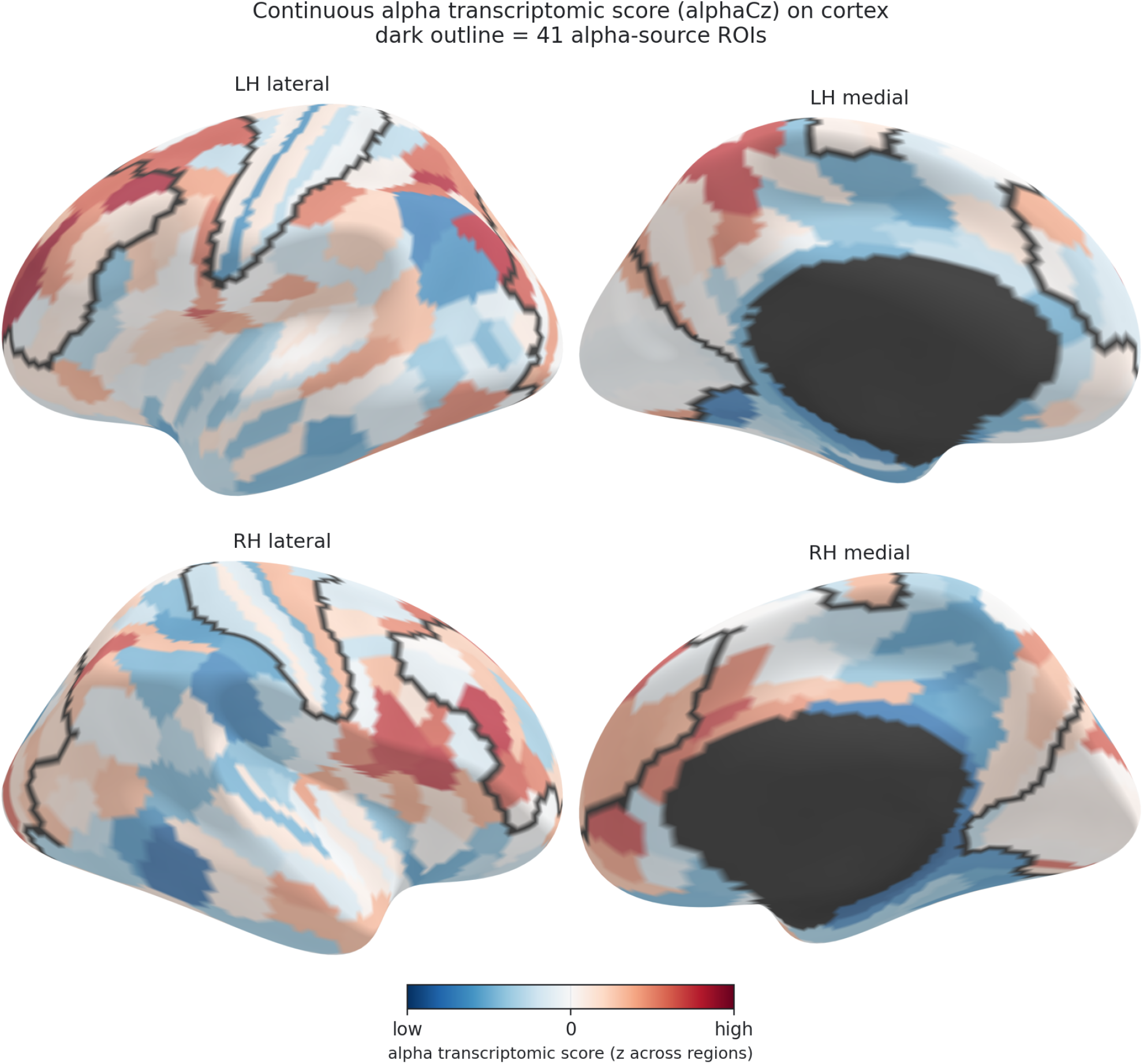
Alpha transcriptomic score on the cortex. The continuous alpha score (alphaCz) per Glasser region, tertile-binned for display, on the inflated surface; the black outline marks the 41 alpha-source ROIs. The score is weakly and diffusely patterned and is not concentrated within the target ROIs.

**Figure 6:**
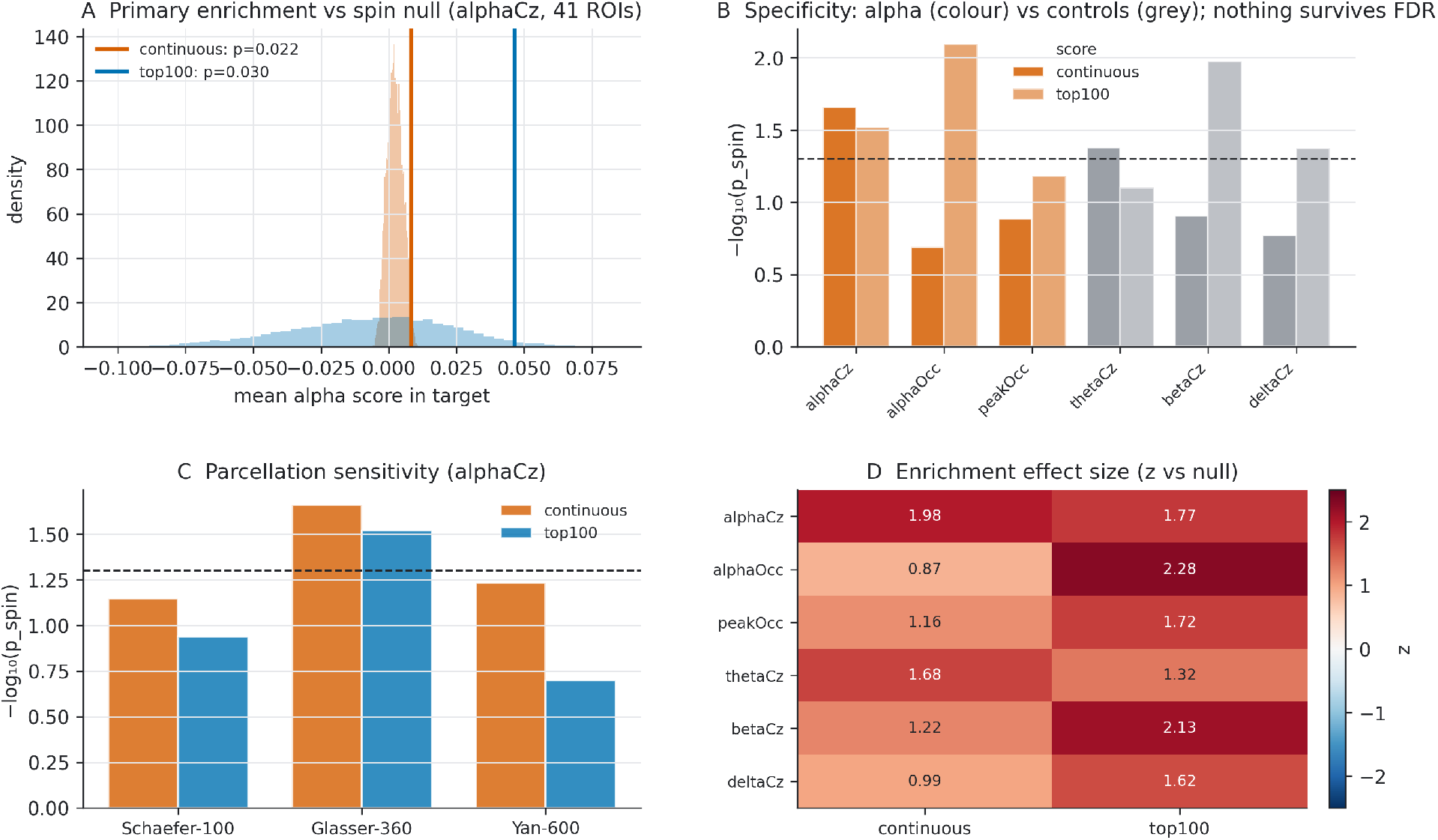
Enrichment, specificity and robustness. (A) Primary alphaCz enrichment against the spin null (both scores). (B) Specificity across the six phenotypes (alpha in colour, controls in grey; dashed line *p* = 0.05); control bands enrich comparably or more, and nothing survives FDR. (C) Parcellation sensitivity: nominal significance only in the native Glasser space. (D) Enrichment effect size (*z* vs null) across phenotypes and scores.

This entirely reframes the result reported in Section 3.3. The spin test asks whether the *brain map* could have arisen by chance given its spatial smoothness, and answers no. It cannot ask whether the *gene set* is special, because it never permutes genes. The apparent enrichment is therefore not evidence about alpha genetics at all. It is a property of how any co-expressed set of genes distributes over this cortical territory, exactly the inflation Fulcher et al. (2021) predict and that a spatial-only null cannot see.

### 3.6 Third test: significance does not survive a change of parcellation

For the primary phenotype, the direction of the continuous-score enrichment was consistent across all three parcellations (positive; *z* between 1.5 and 2.0), but nominal significance was reached only in the native Glasser space (Table 2). Coarsening the parcellation to Schaefer-100, which more closely matches the spatial resolution of MEG source reconstruction, did **not** strengthen the effect (*p*_spin = 0.071); refining it to Yan-600 gave a similar trend (*p*_spin = 0.058). The top-100 score was significant only in the Glasser parcellation. The enrichment therefore does not survive a change of parcellation, and the resolution-matching prediction, that a coarser atlas would recover a signal blurred at fine resolution, was not borne out.

**Table 2.** Parcellation sensitivity for central alpha power (alphaCz). Target counts reflect the fraction of parcels overlapping the Glasser alpha-source set.

| Parcellation | Parcels | Targets | Score | $z$ vs null | $p_{\text{spin}}$ |
| --- | --- | --- | --- | --- | --- |
| Schaefer-100 | 100 | 31 | Continuous | 1.53 | 0.071 |
| Schaefer-100 | 100 | 31 | Top-100 | 1.19 | 0.115 |
| **Glasser-360** | 360 | 82 | **Continuous** | **1.98** | **0.022** |
| **Glasser-360** | 360 | 82 | **Top-100** | **1.77** | **0.030** |
| Yan-600 | 600 | 180 | Continuous | 1.64 | 0.058 |
| Yan-600 | 600 | 180 | Top-100 | 0.94 | 0.200 |

## 4. Discussion

We asked whether the genetic architecture of the EEG alpha rhythm is transcriptomically concentrated in the cortical generators of that rhythm. Across a pre-specified primary test, a specificity panel of three control bands and two further alpha phenotypes, and three parcellations, the answer is negative. A nominally significant enrichment appears for one alpha phenotype in one parcellation, but it is not specific to alpha (control bands enrich comparably or more strongly), it fails to replicate in the other alpha phenotypes, it is indistinguishable from random gene sets, it does not hold across parcellations, and it does not survive multiple-comparison correction. H1 is not supported.

### 4.1 What the null models do and do not establish

The more transferable result concerns H2. Had we followed the common analytic recipe (gene-based statistics, regional expression, spin test), we would have reported a positive finding, and it would have looked solid. Two scores agreed. An independent surrogate model (brainsmash) returned essentially the same *p*-value, which is often taken as evidence of robustness. A positive control confirmed the pipeline was sensitive. And the effect had an obvious mechanistic story. None of that was wrong; it was simply answering a narrower question than it appeared to.

A spin test asks: *given this brain map’s spatial smoothness, could its overlap with the target regions have arisen by chance?* It permutes space, never genes. It therefore cannot detect that the same overlap would have been obtained with almost any gene set of comparable size — which is what the gene-set null revealed (*p*_geneset = 0.33). The two nulls are not redundant strategies for the same problem; they control different confounds, and passing one says nothing about the other. Cross-checking a spin test against another *spatial* null model, as we did with brainsmash, does not help either: two spatial nulls agreeing tells you the spatial correction is stable, not that the gene set is special.

We therefore suggest a minimum standard for enrichment claims in imaging transcriptomics: report a gene-set null alongside the spatial null, and treat a result that passes only the latter as uninterpreted rather than positive. The argument is not new; Fulcher et al. (2021) set it out. What we have not seen is a real case in which a result that passes the spin test, agrees with a surrogate model, and clears a positive control turns out, with the outcome unknown in advance, to be explained entirely by co-expression. The cost of the additional null is negligible: in this study it was a few minutes of computation on results already in hand.

### 4.2 What bounds the substantive null

Four considerations bound the interpretation of the H1 null. First, **statistical power**: the ENIGMA-EEG GWAS, while the largest dedicated EEG-power meta-analysis, is modest by contemporary standards, and only a single gene survived genome-wide correction in our gene-based analysis. Low gene-based signal-to-noise limits the sensitivity of any downstream enrichment, and could in principle mask a true but small effect. Three features of the design bound this concern. Our continuous score deliberately used the *entire* genome-wide gene-level signal rather than a sparse significant set, which is the analysis best suited to a low-powered GWAS; it too was non-specific. The positive control shows the enrichment machinery is not insensitive: it recovered a genuine expression gradient at *p*_spin = 2 × 10^−4^, so the alpha null is not a floor effect of an underpowered *test*, even if the underlying GWAS is underpowered. And the co-expression-aware gene-set null indicates that even the nominal alpha enrichment is a generic property of arbitrary gene sets in this territory (*p*_geneset = 0.33), not a weak-but-real alpha signal awaiting more power.

Second, **the shared genetic architecture of neighbouring EEG bands**. Band-limited EEG powers are strongly genetically correlated (Smit et al., 2018), and the alpha and theta bands in particular share generators and heritable variance. The comparable enrichment we observe for theta, beta and delta is therefore not surprising: to the extent that any oscillatory-power genetics is over-represented in this occipito-parieto-frontal territory, it reflects EEG-power genetics in general rather than an alpha-specific molecular signature. This reframes, rather than rescues, the hypothesis: the alpha-source network is not transcriptomically privileged for alpha genes specifically.

Third, and most consequential for the imaging-transcriptomics framing, **the spatial resolution of the functional target**. The alpha sources were localised from MEG with minimum-norm estimation, whose point-spread is on the order of centimetres; the “alpha-source” label attached to a given fine-grained parcel is therefore spatially uncertain, and the true generators are smeared across neighbouring parcels. This produces a resolution mismatch with *abagen*’s parcel-level expression that biases any localised contrast toward the null. We tested the corollary prediction directly: coarsening the parcellation toward MEG resolution should recover a blurred signal. It did not (Schaefer-100 was, if anything, weaker than Glasser-360), which argues against a large effect merely hidden by resolution mismatch. Switching the inverse operator would not resolve this either. All M/EEG source-reconstruction methods share centimetre-scale point-spread because the inverse problem is ill-posed, and beamformers in particular suppress coherent sources, precisely the phase-coupled parieto-frontal network at issue here, which makes them a poor remedy for this target.

Fourth, **the phenotype itself conflates periodic and aperiodic activity**. ENIGMA-EEG’s band-power phenotypes are computed as the mean spectral power within a fixed frequency window and do not separate the periodic (oscillatory) component from the aperiodic 1/f background, namely the offset and exponent that index cortical excitation/inhibition balance (Gao et al., 2017) and that FOOOF/*specparam* parametrisation was developed to isolate (Donoghue et al., 2020). Because band-limited power is a mixture of both, a fraction of the “alpha power” association signal we carried into MAGMA may in fact index aperiodic, not oscillatory, genetic variation. This is not merely a hypothetical concern: the largest prior EEG GWAS found a genome-wide association between *GABRA2* and beta power (Hodgkinson et al., 2010); since GABAergic signalling is a principal determinant of the aperiodic exponent, this is consistent with band-power phenotypes already carrying an aperiodic signature. If the true signal lies in the aperiodic component, then the band-power test we ran may not just be underpowered for alpha; it may be aimed at a partially different phenotype from the one our oscillatory hypothesis concerns.

#### Strengths

The study is, to our knowledge, the first to bring an EEG-trait GWAS into imaging transcriptomics, and it is deliberately built to resist the over-interpretation that a single positive test invites. The gene-based pipeline is validated by recovery of the expected chr3p21 locus; the target regions are defined natively in the analysis parcellation, avoiding subjective mapping; sensitivity is established by positive control; and specificity, replication, gene-set and parcellation robustness are all examined and reported. Because the additional nulls were specified as part of the design rather than added after an unwelcome result, the reversal in Section 3.5 is a finding rather than a post-hoc rescue.

#### Relevance to brain-state-dependent stimulation

Closed-loop EEG–TMS reads out and targets the same alpha generators, so a molecular signature there would in principle have offered a biological criterion for target selection. Our result gives no support for that: at current GWAS power, cortical alpha-target selection is unlikely to be informed by the regional molecular architecture of alpha-associated genes, and should continue to rest on functional and connectivity criteria. We note this as an implication rather than a demonstration; no stimulation data are analysed here.

## 5. Limitations

Beyond the four interpretive bounds above, several limitations apply. (i) The AHBA comprises six donors, only two with right-hemisphere tissue, and we used five (donor 15496 being unavailable at source); regional expression estimates therefore carry substantial uncertainty, especially for the right hemisphere and for inter-individual variability. (ii) A **modality gap** separates the genetic phenotype from the region set: the GWAS phenotypes are scalp *EEG* band power, whereas the generators were localised from resting-state *MEG*. The two modalities have different sensitivity profiles (MEG is relatively insensitive to radial sources), so the MEG-derived generators are an imperfect proxy for the origins of the EEG signal that was actually genotyped (and, downstream, for the targets of EEG-based closed-loop stimulation). (iii) The alpha-source set is a curated selection of 41 regions chosen to give broad coverage while avoiding proximity-induced spurious connectivity, not an exhaustive generator map; its large frontal contingent may partly reflect connectivity hubs rather than primary generators. (iv) Gene-set enrichment depends on gene-set definition; we mitigated this with a threshold-free continuous score, a top-N sensitivity, and a co-expression-aware gene-set null, but other definitions could differ. (v) Correspondence between regional gene expression and function is correlational and does not license mechanistic claims about TMS-induced plasticity. (vi) The analysis is restricted to cortex and to common variants tagged by the reference panel. (vii) The GWAS phenotypes are band-limited power, which conflates periodic and aperiodic spectral components (see Discussion); no GWAS of aperiodic parameters is currently available to test the two separately.

## 6. Future directions: an aperiodic-component GWAS

The periodic/aperiodic conflation above points to the most direct next step: repeating this pipeline with gene-based statistics derived from a GWAS of the **aperiodic exponent and offset** rather than band power. This alternative phenotype has three properties that make it, on priors, a better-motivated target for imaging transcriptomics than band power. It has a clearer molecular interpretation (excitation/inhibition balance, with GABAergic and glutamatergic receptor genes as natural candidates); resting EEG band power itself is highly heritable (Smit et al., 2005), which, given that band power partly reflects the aperiodic component (see above), is at minimum consistent with a heritable aperiodic contribution, though the aperiodic component’s heritability has not, to our knowledge, been directly estimated by twin or family design; and, unlike band power, it follows a graded sensorimotor-to-association cortical topography that plausibly aligns with the dominant transcriptomic gradient of the AHBA, the same gradient our positive control (Sections 2.6 and 3.2) shows the pipeline can detect.

This analysis is not possible with existing secondary data: to our knowledge no GWAS of FOOOF/*specparam* aperiodic parameters with public summary statistics currently exists. ENIGMA-EEG is a distributed meta-analytic consortium: member cohorts retain their raw EEG and genotypes locally and contribute only summary statistics from an agreed pipeline, so there is no repository of raw EEG one can request; generating this GWAS would mean either (a) proposing an aperiodic-parameter sub-study to the ENIGMA-EEG working group, so that member cohorts run a shared FOOOF pipeline locally and contribute new summary statistics, or (b) a smaller proof-of-concept in a single cohort with both resting EEG and genotypes under controlled access (e.g., COGA via dbGaP), underpowered relative to ENIGMA-EEG but sufficient for an initial test. Either route is a primary-data undertaking beyond the scope of the present secondary-data analysis. The pipeline described here is phenotype-agnostic at the point past MAGMA (Fig. 1) and would consume aperiodic gene-based statistics without modification.

## 7. Conclusion

Genes associated with EEG alpha power show no regionally specific transcriptomic concentration in the cortical generators of the alpha rhythm. Any tendency toward enrichment is weak, shared with the non-alpha bands we tested, fragile to parcellation, and, most importantly, indistinguishable from what arbitrary gene sets produce in the same territory. Establishing whether a true band-specific signal exists will require a substantially larger EEG GWAS, generator maps defined at a spatial resolution commensurate with transcriptomic parcellations, and separation of the periodic and aperiodic components that present band-power phenotypes conflate (Section 6).

The methodological point outlasts this dataset. An enrichment that passed the spin test, agreed with a surrogate model, cleared a positive control, and came with a ready mechanistic interpretation was still explained entirely by gene-set co-expression. Correcting for spatial autocorrelation is a genuine requirement, but it does not settle whether a gene set is special, and adding further *spatial* nulls does not change that. The gene-set null is what settles the question, at a cost of minutes; without it, an unknown fraction of regional enrichment findings in this literature may be properties of the cortex rather than of the genes.

## Data and code availability

Analysis code and derived results (gene-based statistics, region × gene expression summaries, enrichment tables, run logs) are openly available at https://github.com/Jacoposchenetti/Neurogenetics under an MIT licence, and archived at Zenodo: https://doi.org/10.5281/zenodo.21514560. Raw data are not redistributed and must be obtained from their sources under the applicable terms: ENIGMA-EEG summary statistics (ENIGMA-EEG working group), the Allen Human Brain Atlas (via *abagen*), and the 1000 Genomes / MAGMA reference files (CTG lab). All parameters, random seeds and software versions are recorded in the repository run logs, and every figure and reported value can be regenerated from the archived code.

## Author contributions

J.S. is the sole author and was responsible for conceptualisation, methodology, formal analysis, software, visualisation, and writing.

